# Simultaneous stimulation of RuBP regeneration and electron transport increases productivity and water use efficiency under field conditions

**DOI:** 10.1101/2020.01.27.920843

**Authors:** Patricia E. López-Calcagno, Kenny L. Brown, Andrew J. Simkin, Stuart J. Fisk, Tracy Lawson, Christine A. Raines

**Author notes:** Address correspondence to C.A.R. or P.E.L.C.

## Abstract

Previous studies have demonstrated that independent stimulation of either electron transport or RuBP regeneration can increase the rate of photosynthetic carbon assimilation and plant biomass. In this paper, we present evidence that a multi-gene approach to simultaneously manipulate these two processes provides a further stimulation of photosynthesis. We report on the introduction of the cyanobacterial bifunctional enzyme fructose-1,6-bisphosphatase/sedoheptulose-1,7-bisphosphatase or overexpression of the plant enzyme sedoheptulose-1,7-bisphosphatase, together with expression of the red algal protein cytochrome *c*_6_, and show that a further increase in biomass accumulation under both glasshouse and field conditions can be achieved. Furthermore, we provide evidence that the simultaneous stimulation of electron transport and RuBP regeneration can lead to enhanced intrinsic water use efficiency under field conditions.

**One sentence summary:** Simultaneous stimulation of RuBP regeneration and electron transport results in improvements in biomass yield in glasshouse and field grown tobacco.

---

Yield potential of seed crops grown under optimal management practices, and in the absence of biotic and abiotic stress, is determined by incident solar radiation over the growing season, the efficiency of light interception, energy conversion efficiency and partitioning or harvest index. The “harvest index” is the amount of total energy partitioned into the harvestable portion of the crop, whilst the “conversion efficiency” is the ratio of biomass energy produced over a given period divided by light energy intercepted by the canopy over the same period^1^. For the major crops, the only component not close to the theoretical maximum is energy conversion efficiency, which is determined by gross canopy photosynthesis minus respiration. This highlights photosynthesis as a target for improvement to raise yield potential in major seed crops^1^.

Transgenic experiments and modelling studies have provided compelling evidence that increasing the levels of photosynthetic enzymes in the Calvin Benson (C_B_) cycle has the potential to impact photosynthetic rate and yield^1–14^. Early experimental evidence illustrated that even small reductions in the CB enzymes sedoheptulose-1,7-bisphosphatase (SBPase^15–17^), fructose-1,6-bisphophate aldolase (FBPA^18,19^) and the chloroplastic fructose-1,6-bisphosphatase (FBPase^20–22^) negatively impact on carbon assimilation and plant growth. These studies indicated that these enzymes exercise significant control over photosynthetic efficiency, suggesting that improvements in photosynthetic carbon fixation may be achieved and maintained through manipulation of CB cycle enzymes^11, 23^. These results also demonstrated that, although some enzymes exert more control over the CB cycle than others, there is no single limiting step in photosynthetic carbon assimilation. Over the last two decades a number of transgenic studies have supported this. Over-expression of SBPase in tobacco^3,5,6^, Arabidopsis^7^, tomato^13^ and wheat^24^ has demonstrated the potential of manipulating the expression of CB cycle enzymes and specifically the regeneration of RuBP to increase growth, biomass (30-42%) and even seed yield (10-53%). Similarly, overexpression of other enzymes including FBPA^12^, cyanobacterial SBPase, FBPase^25^ and the bifunctional fructose-1,6-bisphosphatases/sedoheptulose-1,7-bisphosphatase (FBP/SBPase^2,26,27^) in a range of species including tobacco, lettuce and soybean has shown that increasing photosynthesis increases yield, reinforcing the original hypothesis that manipulating the activity of the CB cycle enzymes can be used to increase productivity.

In addition to manipulation of CB cycle genes, increasing photosynthetic electron transport has also been shown to have a beneficial effect on plant growth. Overexpression of the Rieske FeS protein -a key component of the cytochrome *b_6_f* complex-in Arabidopsis, has previously been shown to lead to increases in electron transport rates, CO_2_ assimilation, biomass and seed yield^28^. Similarly, the introduction of the algal cytochrome *c_6_* protein into Arabidopsis and tobacco^29,30^ resulted in increased growth. In these transgenic plants, electron transport rate was increased along with ATP, NADPH, chlorophyll, starch content, and capacity for CO_2_ assimilation. Higher plants have been proposed to have lost the cytochrome *c_6_* protein through evolution, but in green algae and cyanobacteria, which have genes for both cytochrome *c_6_* and plastocyanin (PC), cytochrome *c_6_* has been shown to replace PC as the electron transporter connecting the cytochrome *b_6_/f* complex with PSI under Cu deficiency conditions^31,32^. There is evidence showing that PC can limit electron transfer between cytochrome *b_6_f* complex and PSI^33^, and in Arabidopsis, it has been shown that introduced algal cytochrome *c_6_* is a more efficient electron donor to P700 than PC^29^. This evidence suggests the introduction of the cytochrome *c_6_* protein in higher plants as a viable strategy for improving photosynthesis.

Previous research has shown that taking a multi-gene approach to increase the levels of more than one enzyme or protein simultaneously can result in a cumulative increase in photosynthesis and biomass yield^6,7,34^. Building on this approach, the work in this paper aims to test the hypothesis that combining an increase in the activity of a CB cycle enzyme, specifically enhancing RuBP regeneration, together with stimulation of the electron transport chain can boost photosynthesis and yield above that observed when these processes are targeted individually. To test this hypothesis *Nicotiana tabacum* plants expressing the cyanobacterial FBP/SBPase or the higher plant SBPase, and the algal cytochrome *c*_6_ were generated in two different tobacco cultivars. The analysis presented here demonstrates that the simultaneous stimulation of electron transport and RuBP regeneration leads to a significant increase in photosynthetic carbon assimilation, and results in increased biomass and yield under both glasshouse and field conditions.

## RESULTS

### Production and Selection of Tobacco Transformants

Previous differences observed in the biomass accumulation between Arabidopsis and tobacco overexpressing SBPase and SBPase plus FBPA^6,7^ led us to explore the effect of similar manipulations (RuBP regeneration by overexpression of SBPase or introduction of the cyanobacterial FBP/SBPase, together with enhanced electron transport) in on two different tobacco cultivars with very different growth habits: *N. tabacum* cv. Petite Havana and *N. tabacum* cv. Samsun. Sixty lines of cv. Petit Havana, and up to fourteen lines of cv. Samsun were generated per construct and T0 and T1 transgenic tobacco were screened by qPCR analysis to select independent lines with expression of the transgenes (data not shown).

*N. tabacum* cv. Petit Havana T2/T3 progeny expressing FBP/SBPase (S_B_; lines S_B_03, S_B_06, S_B_21, S_B_44) or cytochrome *c*_6_ (C_6_; lines C15, C41, C47, C50) and cv. Samsun lines expressing SBPase + cytochrome *c*_6_ (SC_6_, lines 1, 2 and 3) were produced by agrobacterium transformation. *N. tabacum* cv. Petit Havana plants expressing both S_B_ and C_6_ were generated by crossing S_B_ lines (06, 44, 21) with C_6_ lines (15, 47, 50) to generate four independent S_B_C_6_ lines: S_B_C1 (S_B_06xC47), S_B_C2 (S_B_06xC50), S_B_C3 (S_B_44xC47) and S_B_C6 (S_B_21xC15). Semi-quantitative RT-PCR was used to detect the presence of the FBP/SBPase transcript in lines S_B_ and S_B_C_6_, cytochrome *c*_6_ in lines C_6_, S_B_C_6_ and SC_6_, and SBPase in lines S and SC_6_ (**Fig. 1a**). We analysed total extractable FBPase activity in the leaves of the cv. Petite Havana T2/T3 & F3 homozygous progeny lines used to determine chlorophyll fluorescence and photosynthetic parameters. This analysis showed that these plants (S_B_ and S_B_C_6_) had increased levels of FBPase activity ranging from 34 to 47% more than the control plants (**Fig. 1b**). The S and SC_6_ lines were from the same generation of transgenic plants used in a previous study and shown to have increased SBPase activity^6^.

**Fig 1.**
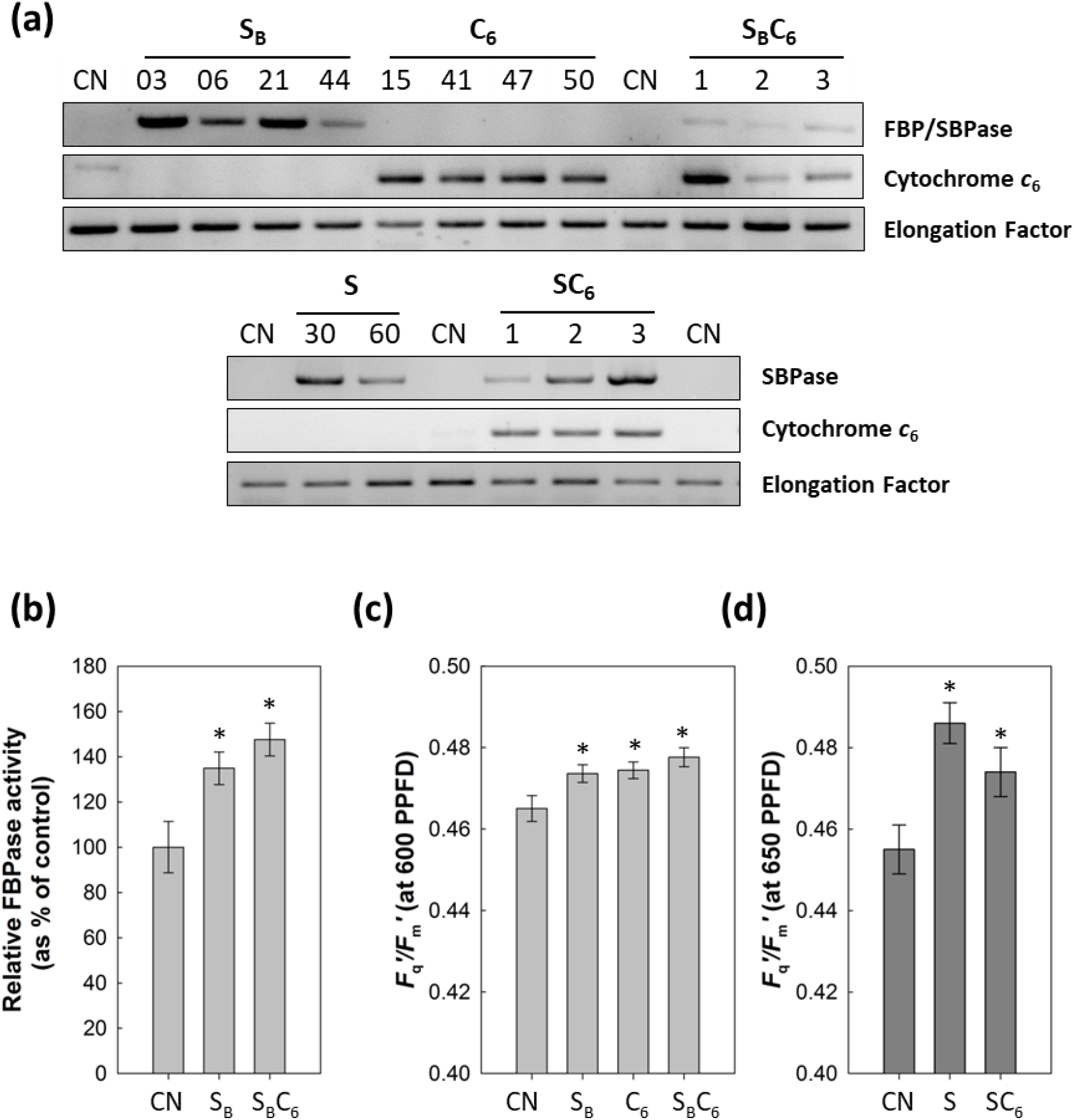
Screening of transgenic plants overexpressing FBP/SBPase, SBPase, and cytochrome *c*_6_. (**a**) Transcript levels in S_B_, C_6_, S_B_C_6_, S and SC_6_ lines compared to controls (CN). (**b**) FBPase activity in S_B_ and S_B_C_6_ lines relative to controls. (**c-d**) Chlorophyll fluorescence imaging of plants grown in controlled environmental conditions used to determine *F*_q_’/*F*_m_’ (maximum PSII operating efficiency) at 600-650 μmol m^−2^ s^−1^. n=7-10. * indicates lines which are statistically different to control groups (p<0.05).

Chlorophyll fluorescence analysis confirmed that in young plants, the operating efficiency of photosystem two (PSII) photochemistry *F*_q_’/*F*_m_’ at an irradiance of 600-650 μmol m^−2^ s^−1^ was significantly higher in all selected lines compared to either WT or null segregant controls (**Fig. 1c, d**). However, the *F*_q_’/*F*_m_’ values of the S_B_C_6_ and SC_6_ lines, were not significantly different from the *F*_q_’/*F*_m_’ values obtained from the plants expressing individually FBP/SBPase (S_B_), cytochrome *c*_6_ (C_6_) or SBPase (S).

### Stimulation of electron transport and RuBP regeneration increases photosynthetic performance in two distinct tobacco varieties under glasshouse conditions

Transgenic lines selected based on the initial screens described above were grown in the glasshouse, in natural light supplemented to provide illumination between 400-1000 μmol m^−2^ s^−1^. The rate of net CO_2_ assimilation (*A*) and *F*_q_’/*F*_m_’ was determined as a function of internal CO_2_ concentration (*C*_i_), in mature and developing leaves of *N. tabacum* cv. Samsun (S and SC_6_) and in mature leaves of *N. tabacum* cv. Petit Havana (S_B_, C_6_ and S_B_C_6_) (**Fig. 2a**). The transgenic lines displayed greater CO_2_ assimilation rates than that of the control (CN) plants. *A* was 15% higher than the controls in the mature leaves of the SC_6_, at a *C*_i_ of approximately 300 μmol mol^−1^ (equivalent to current atmospheric [CO_2_]) (**Fig. 2b**). The developing leaves of the SC_6_ plants also showed significant increases in PSII operating efficiency (*F*_q_’/*F*_m_’) and in the PSII efficiency factor (*F*_q_’/*F*_v_’; which is determined by the ability of the photosynthetic apparatus to maintain Q_A_ in the oxidized state and therefore a measure of photochemical quenching) when compared to control plants (**Fig. 2c**). Interestingly, in mature leaves of the cv. Samsun transgenic plants, the differences in assimilation rates and in the operating efficiency of PSII photochemistry between the transgenic and the CN plants were smaller than in the developing leaves. Only the S transgenic plants displayed a higher average value for *F*_q_’/*F*_m_’ and *F*_q_’/*F*_v_’ than the CN plants at all CO_2_ concentrations measured. In contrast, the mature leaves of SC_6_ plants displayed *F*_q_’/*F*_v_’ values higher than the control only at *C*_i_ levels between 300 and 900 μmol mol^−1^ (**Fig. 2b**).

**Fig 2.**
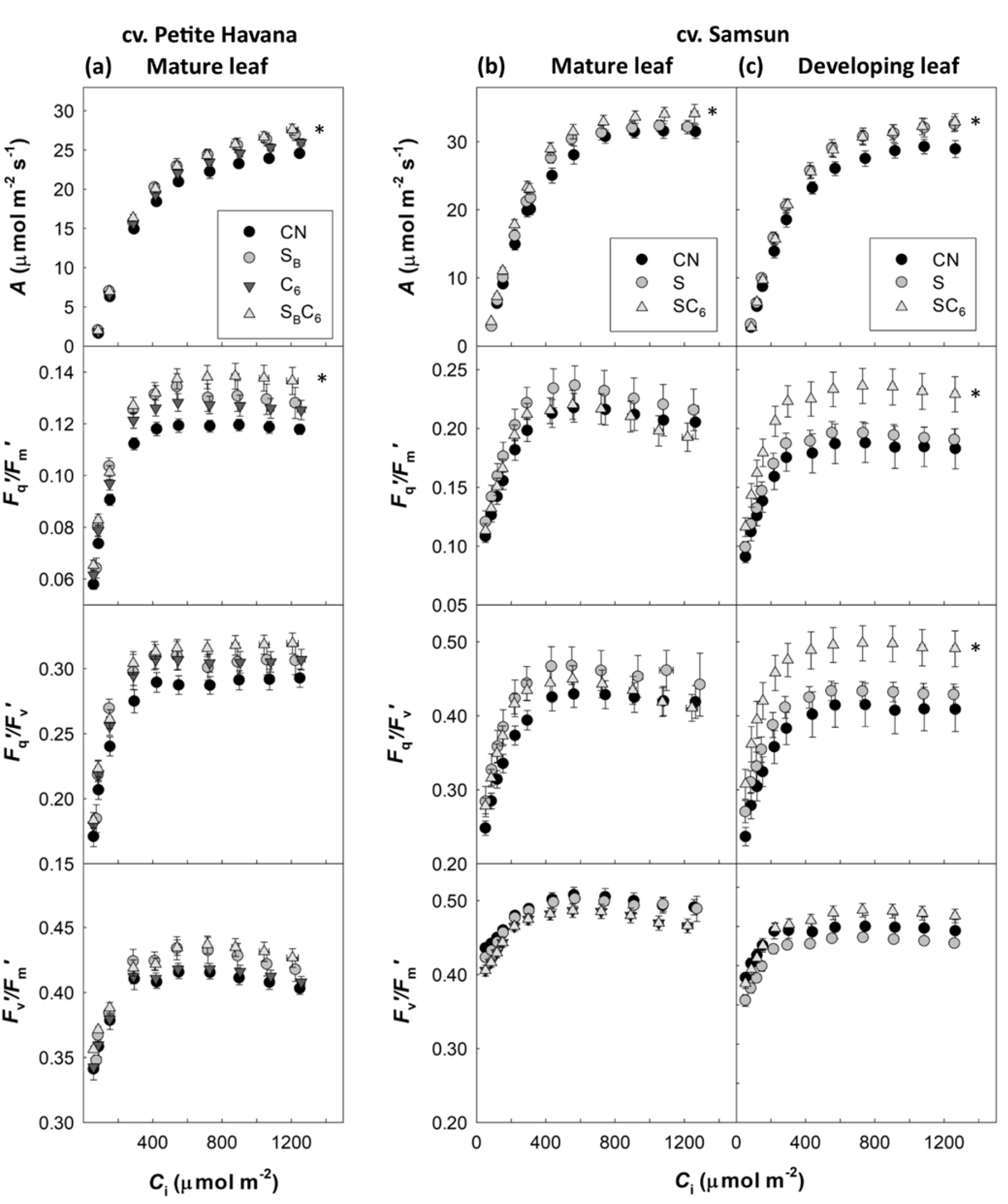
Photosynthetic responses of transgenic plants grown in glasshouse. Photosynthetic carbon fixation rates, actual operating efficiency of PSII in the light (*F*_q_’/*F*_m_’), electron sinks pulling away from PSII (*F*_q_’/*F*_v_’) and PSII maximum efficiency (*F*_v_’/*F*_m_’) are presented in (**a**) mature leaves of cv. Petit Havana and (**b**) mature and (**c**) developing leaves of cv. Samsun. Parameters were determined as a function of increasing CO_2_ concentrations at saturating-light levels in developing (11-13cm in length) and mature leaves from control and transgenic plants. Plants were grown in natural light conditions in the glasshouse where light levels oscillated between 400 and 1000 μmol m^−2^ s^−1^ (supplemental light maintain a minimum of 400 μmol m^−2^ s^−1^). Lines expressing FBP/SBPase (S_B_), Cytochrome *c*_6_ (C_6_), FBP/SBPase and Cytochrome *c*_6_ (S_B_C_6_), SBPase (S) and SBPase and Cytochrome *c*_6_ (SC_6_). Control group (CN) represent both WT and azygous plants. Evaluations are based on 3-4 plants individual plants per line, and 3 to 4 independent transgenic lines per manipulation. Asterisks indicate significance between transgenics and control group, using a linear mixed-effects model and type III ANOVA and contrast analysis, *p < 0.05.

Similar trends were shown for the *N. tabacum* cv. Petit Havana transgenic plants, which displayed higher average values of *A*, *F*_q_’/*F*_m_’ and *F*_q_’/*F*_v_’ than the CN (**Fig. 2a**). In the leaves of the S_B_C_6_ plants (cv. Petit Havana) these significant increases were similar to the developing leaves of the SC_6_ lines (cv. Samsun).

The developing leaves of both the S and SC_6_ plants (cv. Samsun) showed a significant increase in *J*_max_ and *A*_max_ when compared the control plants (**Table 1**). The mature leaves of the SC_6_ transgenics also displayed a significantly higher *Vc*_max_, *J*_max_ and *A*_max_ than the CN. In contrast, the leaves of the S_B_C_6_ plants (cv. Petite Havana) only had significant increases in *A*_max_, although higher average values for *Vc*_max_, and *J*_max_ were evident. These results showed that simultaneous stimulation of electron transport and RuBP regeneration by expression of cytochrome *c_6_* in combination with FBP/SBPase or SBPase has a greater impact on photosynthesis than the single manipulations in all plants analysed.

**Table 1.**
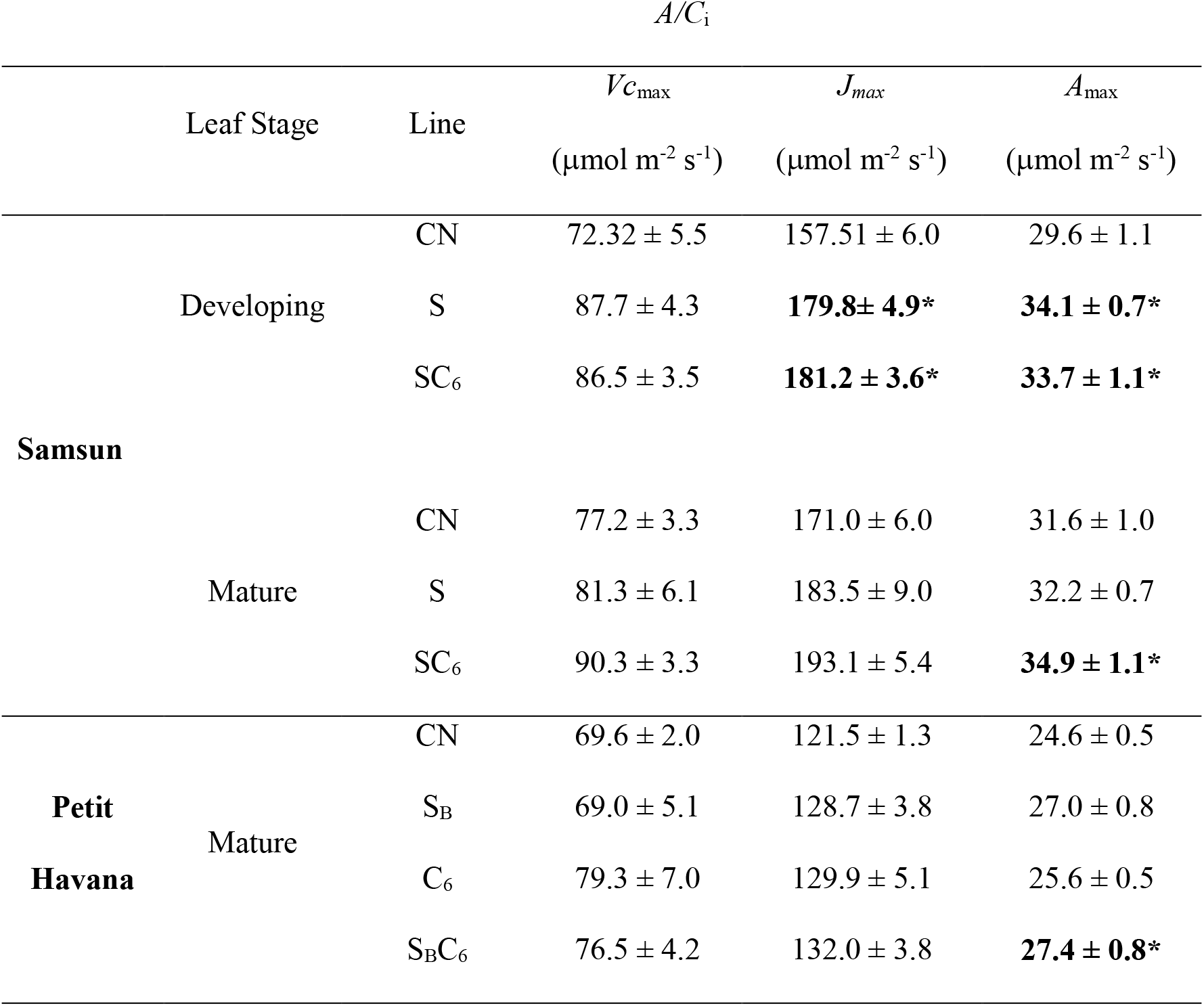
Maximum electron transport rate and RuBP regeneration (*J*_max_), carboxylation rate of Rubisco (*Vc*_max_) and maximum assimilation (*A*_max_) of WT and transgenic lines. Results were determined from the *A*/*C*_i_ curves in Figure 2 using the equations published by von Caemmerer and Farquhar^58^. Statistical differences are shown in boldface (* p<0.05). Mean and SE are shown.

### Stimulation of electron transport and RuBP regeneration stimulates growth in two distinct tobacco varieties under glasshouse conditions

In parallel experiments, plants expressing FBP/SBPase (S_B_), cytochrome *c*_6_ (C_6_) and both (S_B_C_6_) (*N. tabacum* cv. Petite Havana) and plants expressing SBPase (S) and SBPase + cytochrome *c*_6_(SC_6_) (*N. tabacum* cv. Samsun) were grown in the glasshouse for four and six weeks respectively before harvesting. Height, leaf number, total leaf area and above ground biomass were determined (**Fig 3**). All of the transgenic plants analysed here displayed increased height when compared to CN plants. Plants expressing cytochrome *c*_6_(C_6_, S_B_C_6_, (cv. Petite Havana) and SC_6_ (cv. Samsun)) had a significant increase in leaf area and in biomass compared to their respective controls. Notably the S_B_C_6_ and SC_6_ transgenics displayed significantly greater leaf area than the single S_B_ and S transgenic plants respectively. The total increase in above ground biomass when compared to CN group was 35% in S_B_, 44% in C_6_ and 9% in S, with consistently higher means in the double manipulations S_B_C_6_ (52%) and SC_6_ (32%).

**Fig 3.**
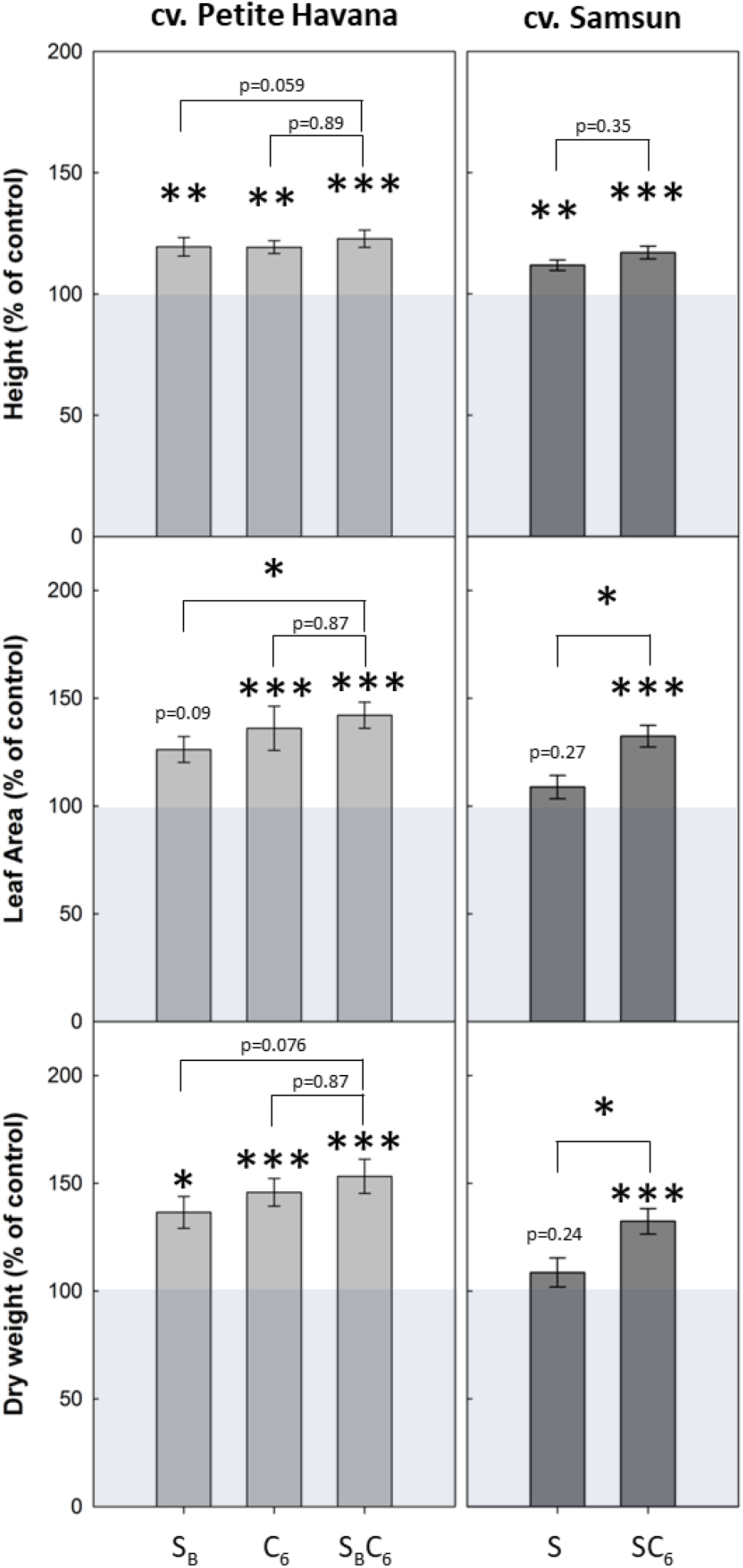
Increased SBPase or expression of FBP/SBPase and cytochrome *c*_6_ increases biomass in glasshouse grown plants. Tobacco plants were germinated in growth cabinets and moved to the glasshouse at 10-14 d post-germination forty-day-old (cv. Petit Havana) or fifty-six-day-old (cv. Samsun) plants were harvested and plant height, leaf area and above-ground biomass (dry weight) were determined. Control group represent both WT and azygous plants. Mean and SE presented. n= 5-6 individual plants from 2 to 4 independent transgenic lines. Asterisks indicate significance between transgenics and control group, or between genotypes using ANOVA with Tukey’s post hoc test, *p < 0.05, **p < 0.01, ***p < 0.001.

### Simultaneous expression of FBP/SBPase and cytochrome *c*_6_ increases growth and water use efficiency under field conditions

To test whether the increases in biomass observed in these transgenic plants under glasshouse conditions could be reproduced in a field environment, a subset of lines was selected for testing in the field. Since the larger percentage increases in biomass were displayed by the manipulations in *N. tabacum* cv. Petit Havana, these plants were selected and tested in three field experiments in two different years (2016 and 2017).

In 2016, a small-scale replicated control experiment was carried out to evaluate vegetative growth in the field, in the lines expressing single gene constructs for FBP/SBPase (S_B_) and cytochrome *c*_6_ (C_6_). Plants were germinated and grown under controlled environment conditions for 25 d before being moved to the field. After 14 d in the field, plants were harvested at an early vegetative stage and plant height, total leaf area and above ground biomass were measured (**Fig 4a**). These data revealed that the S_B_ and C_6_ plants showed an increase in height, leaf area and above ground biomass of 27%, 35% and 25% respectively for S_B_ and 50%, 41% and 36% respectively for C_6_ when compared to CN plants.

**Fig 4.**
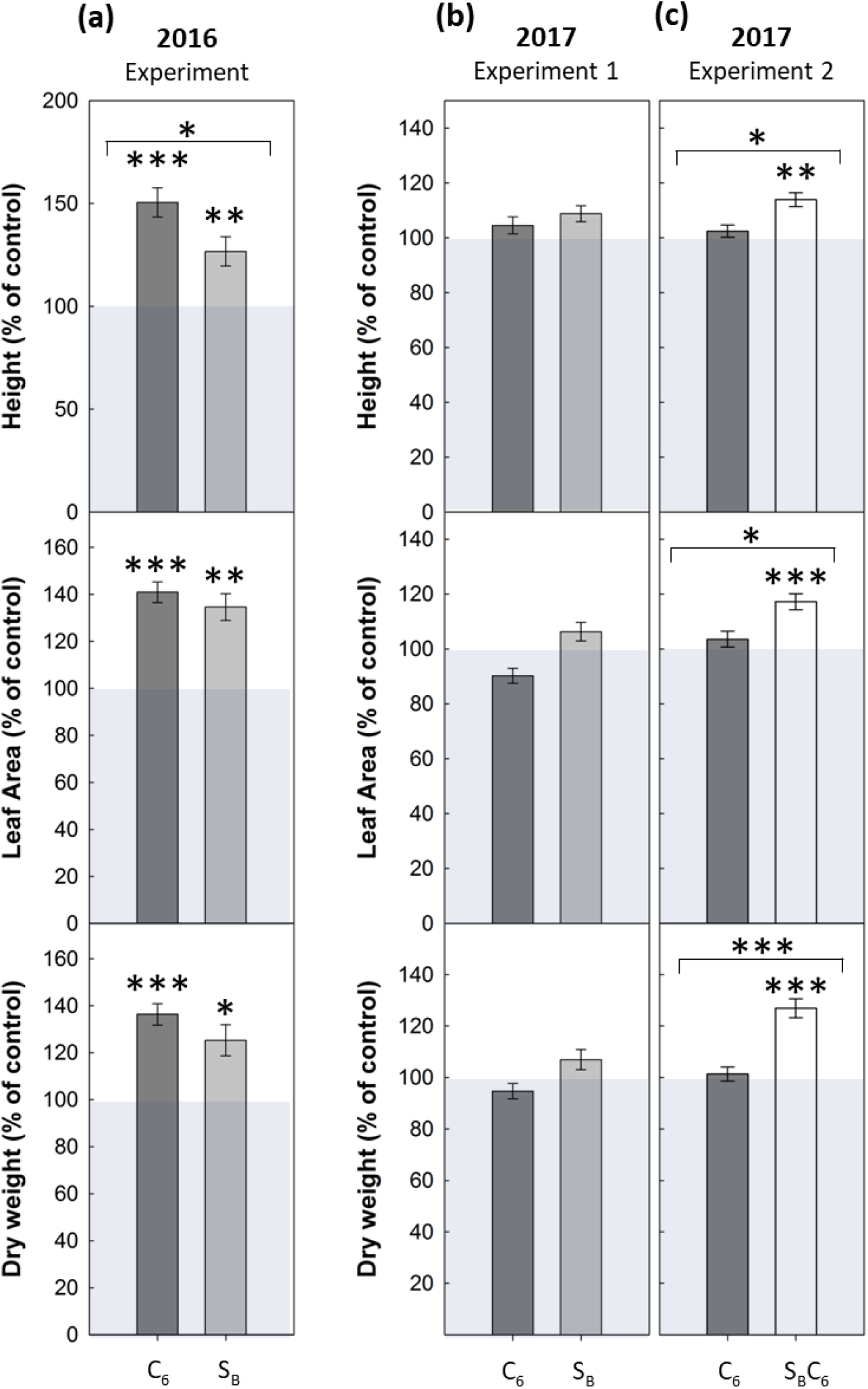
Simultaneous expression of FBP/SBPase and cytochrome *c*_6_ increases biomass in field grown plants. (**a**) Forty-day-old (young) 2016 field-grown plants (plants were germinated and grown in glasshouse conditions for 26 d and then allowed to grow in the field in summer 2016 for 14 d); (**b-c**) Fifty-seven-day-old or sixty-one-day-old (flowering) 2017 field-grown plants (plants were germinated and grown in glasshouse conditions for 26 d and then allowed to grow in the field in summer 2017 until when flowering established, circa 30 d). Light grey bars represent FBP/SBPase expressing plants (S_B_), dark grey bars represent cytochrome *c*_6_ expressing plants (C_6_) and white bars represent plant expressing both transgenes (S_B_C_6_). Plant height, leaf area and total above-ground biomass (dry weight) are displayed. Control group represent both WT and azygous plants. Mean ± SE presented. 2-3 independent lines per manipulation 6 (**a**) or 24 (**b-c**) plants per line. Asterisks indicate significance between transgenics and control group, or between transgenic groups, *p < 0.05, **p < 0.01, ***p < 0.001.

In 2017, two larger scale, randomized block design field experiments were carried out to evaluate performance in the S_B_, C_6_ and S_B_C_6_ plants compared to CN plants. Plants were grown from seed in the glasshouse for 33 d, and then moved to the field and allowed to grow until the onset of flowering (further 24-33 d), before harvesting. In **Fig 4b, c** it can be seen that the S_B_ and C_6_ plants harvested after the onset of flowering did not display any significant increases in height, leaf area or biomass. Interestingly, plants expressing both FBP/SBPase and cytochrome *c*_6_ (S_B_C_6_), displayed a significant increase in a number of growth parameters; with 13%, 17% and 27% increases in height, leaf area and above ground biomass respectively when compared to controls.

Additionally, in the 2017 field experiments *A* as a function of C_*i*_ at saturating light (*A*/*C*_i_) was determined. In the 2017 experiment 1 (Exp.1) a significant increase in *A* was observed in S_B_ and C_6_ plants without differences in PSII operating efficiency (*F*_q_’/*F*_m_’) (**Fig. 5a**). However, in the 2017 experiment 2 (Exp.2), no differences in *A* or in *F*_q_’/*F*_m_’ values were evident in the C_6_ and S_B_C_6_ plants when compared to the CN plants (**Fig. 5b**). Analysis of *A* as a function of light (*A*/*Q*) showed either small or no significant differences in *A* between genotypes (**Fig. 6a**). Interestingly, *g_s_* in the S_B_C_6_ plants were significantly lower than C_6_ and CN plants at light intensities above 1000 μmol m^−2^ s^−1^ (**Fig 6b**), which resulted in a significant increase in intrinsic water use efficiency (*iWUE*) for S_B_C_6_ plants (**Fig 6d**). No significant differences in *iWUE* were observed for S_B_ or C_6_ transgenic plants.

**Fig 5.**
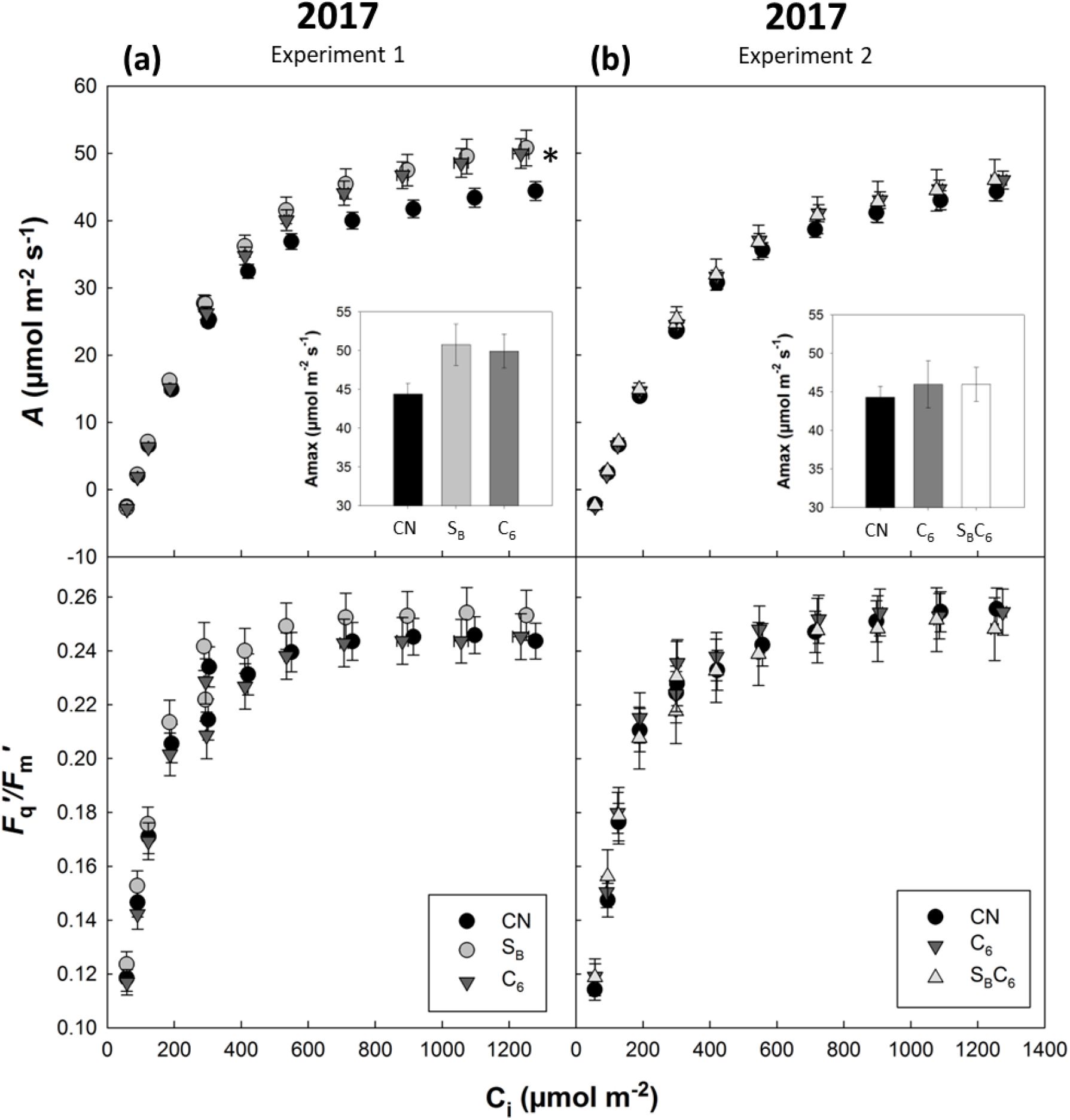
Photosynthetic capacity of field-grown transgenic plants. Photosynthetic carbon fixation rates and operating efficiency of PSII as a function of increasing CO_2_ concentrations at saturating-light levels in mature leaves from CN and transgenic plants. (**a**) 2017 experiment 1: Lines expressing FBP/SBPase (S_B_) and cytochrome *c*_6_ (C_6_). (**b**) 2017 experiment 2: Lines expressing cytochrome *c*_6_ (C_6_) and FBP/SBPase and cytochrome *c*_6_ (S_B_C_6_). Control group (CN) represent both WT and azygous plants. Evaluations are based on 4-5 individual plants from 2-3 independent transgenic lines. Asterisks indicate significance between transgenics and control group, using a linear mixed-effects model and type III ANOVA, *p < 0.05.

**Fig 6.**
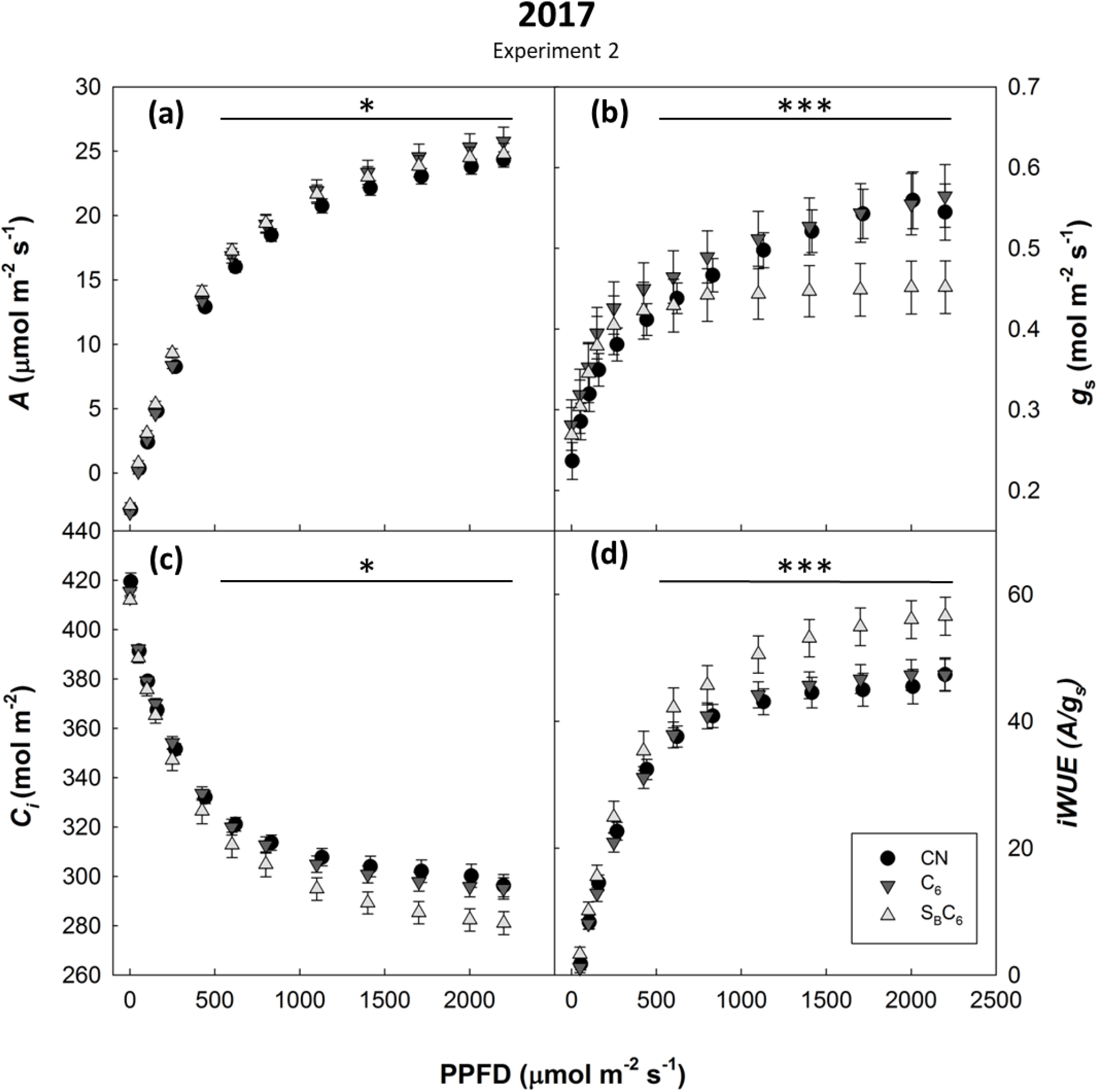
Simultaneous expression of FBP/SBPase and cytochrome *c_6_* can increase water use efficiency under field conditions. (**a**) Net CO_2_ assimilation rate (*A*), (**b**) Stomatal conductance (*g_s_*), (**c**) Intercellular CO_2_ concentration (*C_i_*), and (**d**) Intrinsic water-use efficiency (i*WUE*) as a function of light (PPFD) in field-grown plants. Lines expressing cytochrome *c*_6_ (C_6_) and FBP/SBPase and cytochrome *c*_6_ (S_B_C_6_). Control group (CN) represent both WT and azygous plants. Mean ± SE presented. Evaluations are based on 4-5 individual plants from 2-3 independent transgenic lines per manipulation. Asterisks indicate significance between transgenics and control group, using a linear mixed-effects model and type III ANOVA and contrast analysis, *p < 0.05, **p < 0.01, ***p < 0.001.

## DISCUSSION

In this study, we describe the generation and analysis of transgenic plants with simultaneous increases in electron transport and improved capacity for RuBP regeneration, in two different tobacco cultivars. Here we have shown how independent stimulation of electron transport (by expression of cytochrome *c*_6_) and stimulation of RuBP regeneration (by expression of FBP/SBPase or overexpression of SBPase) increased photosynthesis and biomass in glasshouse studies. Furthermore, we demonstrated how the targeting of these two processes simultaneously (in the S_B_C_6_ and SC_6_ plants) had an even greater effect in stimulating photosynthesis and growth. Additionally, in field studies we demonstrate that plants with simultaneous stimulation of electron transport and of RuBP regeneration had increased *iWUE* with an increase in biomass.

Under glasshouse conditions, increases in photosynthesis were observed in all of the transgenic plants analysed here and these were found to be consistently correlated with increases in biomass. Although increases in photosynthesis and biomass have been reported for plants with stimulation of RuBP regeneration in both model^2,3,6,7,34^ and crop^26,24^ species; and electron transport in Arabidopsis and tobacco^28–30^, the data presented here provides the first report of increased photosynthesis and biomass by the simultaneous stimulation of electron transport and RuBP regeneration. Increases in *A* were observed under glasshouse conditions in the leaves of all of the different transgenic tobacco plants and in both of the tobacco cultivars (cv. Petit Havana and cv. Samsun). Analysis of the *A*/*C*_i_ response curves showed that the average values for the photosynthetic parameters *Vc*_max_, *J*_max_ and *A*_max_ increased by up to 17, 14 and 12% respectively. These results indicated that not only was the maximal rate of electron transport and RuBP regeneration increased, but the rate of carboxylation by Rubisco was also increased. Although this may seem counterintuitive in that we have not targeted directly Rubisco activity, it is in keeping with a study by Wullschleger^35^ of over 100 plant species that showed a linear correlation between *J*_max_ and *Vc*_max_. Furthermore, it has also been shown previously that overexpression of SBPase leads not only to a significant increase in *J*_max_ but that an increase in *Vc*_max_ and Rubisco activation state^3,6^.

Notably, in the greenhouse study, the highest photosynthetic rates were obtained from the leaves of plants in which both electron transport and RuBP regeneration (S_B_C_6_ and SC_6_) were boosted, suggesting that the co-expression of these genes results in an additive effect on improving photosynthesis. In addition to the increases in *A*, the plants with simultaneous stimulation of electron transport and RuBP regeneration displayed a significant increase in *F*_q_’/*F*_m_’, indicating a higher quantum yield of linear electron flux through PSII compared to the control plants. These results are in keeping with the published data for the introduction of cytochrome *c*_6_ and the overexpression of the Rieske FeS protein in Arabidopsis^28,29^. In both of these studies the plants had a higher quantum yield of PSII and a more oxidised plastoquinone pool^29^, suggesting that, although PC is not always limiting under all growth conditions^36^, there is scope to stimulate reduction of PSI by using alternative, more efficient electron donors to PSI^29,33^. Furthermore, in the S_B_C_6_ and SC_6_ plants the increase in *F*_q_’/*F*_m_’ was found to be largely driven by the increase in the PSII efficiency factor (*F*_q_’/*F*_v_’). This suggests that the increase in efficiency in these plants is likely due to stimulation of processes down stream of PSII such as CO_2_ assimilation.

To provide further evidence of the applicability of targeting both electron transport and RuBP regeneration to improve crop yields, we tested these plants in the field. Here we showed that the expression FBP/SBPase alone led to an increase in growth and biomass in the 2016 field-grown plants of between 22-40%, when harvested during early vegetative growth, prior to the onset of flowing. Interestingly, when plants with the same transgenic manipulations were harvested later in development, after the onset of flowering, in the 2017 field trials, this advantage was no longer evident and the single FBP/SBPase expressors were indistinguishable from the control plants. These results are in contrast to the 2016 field data and may be due to the later timing in development of the harvest in the 2017 experiment.

The transgenic plants expressing cytochrome *c*_6_ alone also showed enhanced growth and biomass early development, but as with the FBPase/SBPase plants, this improvement was no longer evident when plants were harvested after flowering. This difference in biomass gain between the early and late harvest was not observed in a parallel experiment, where the overexpression of H-protein was shown to increase biomass under field conditions in plants harvested in early development and after the onset of flowering^37^. These results suggest that the expression of FBP/SBPase or cytochrome *c*_6_ alone, may provide an advantage under particular sets of conditions or at specific stages of plant development. This might be exploitable for some crops where an early harvest is desirable (eg. some types of lettuce, spinach and tender greens)^26^. In contrast with the results with the single manipulations described above, plants simultaneously expressing both cytochrome *c*_6_ and FBP/SBPase displayed a consistent increase in biomass after flowering under field conditions.

In the transgenic lines grown in the field, the correlation between increases in photosynthesis and increased biomass were less consistent than that observed under glasshouse conditions. The significant increases in photosynthetic capacity displayed by the FBP/SBPase and cytochrome *c*_6_ expressors in 2017 Exp. 1, provided clear evidence that these individual manipulations are able to significantly stimulate photosynthetic performance under field conditions. However, no increase in biomass was evident in these plants. In contrast in the 2017 Exp. 2 we did not detect any significant differences in photosynthetic capacity in either the cytochrome *c*_6_ expressors or the plants with simultaneous expression of FBP/SBPase + cytochrome *c*_6_, but increased biomass was evident. At this point we have no explanation for this disparity. However, although not significantly different, in all experiments, the mean *A* values of the transgenic plants were consistently higher than those of the controls. It is known that even small increases in assimilation throughout the lifetime of a plant will have a cumulative effect, which could translate into a significant biomass accumulation^6^, this may in part explain the disparity with the biomass results presented.

An unexpected result that was found only in the plants with simultaneous expression of FBP/SBPase + cytochrome *c*_6_ (S_B_C_6_), was that they had a lower *g*_s_ and lower *C*_i_ concentration at light intensities above 1000 μmol m^−2^ s^−1^, when compared to control plants. Normally lower *C*_i_ would be expected to lead to a reduction in photosynthesis but, interestingly, these plants were able to maintain CO_2_ assimilation rates equal to or higher than control plants resulting in an improvement in *iWUE*. A similar improvement in *iWUE* was seen in plants overexpressing the NPQ related protein, PsbS^38^. It was shown that light-induced stomatal opening was reduced in these plants which had a more oxidized Q_A_ pool, which has been proposed to act as a signal in stomatal movement^39^. Our results provide further support for the proposal that the increased capacity for photosynthesis in the S_B_C_6_ plants is compensating for the reduction in *C*_i_. This higher *iWUE* and the fact that a higher productivity than controls has been reported in field studies for transgenic lines with increased RuBP regeneration grown under CO_2_ enrichment^5,26^, highlight the potential of manipulating electron transport and RUBP regeneration in the development of new varieties able to sustain photosynthesis and yields under climate change scenarios.

## MATERIALS AND METHODS

### Generation of constructs and transgenic plants

Constructs were generated using Golden Gate cloning^40,41^ or Gateway cloning technology^42^. Transgenes were under the control of CaMV35S and FMV constitutive promoters.

For *N. tabacum* cv. Petit Havana, the codon optimised cyanobacterial bifunctional fructose-1,6-bisphosphatases/sedoheptulose-1,7-bisphosphatase (FBP/SBPase; *slr2094* Synechocystis sp. PCC 7942^2^ linked to the geraniol synthase transit peptide^43^ and the codon optimised *P. umbilicalis’s* cytochrome *c*_6_ (AFC39870) with the chlorophyll a-b binding protein 6 transit peptide from Arabidopsis (AT3G54890) were used to generate Golden Gate^41^ over-expression constructs (EC23083 and EC23028) driven by the FMV^44^ and CaMV 35S promoters respectively.

For *N. tabacum* cv. Samsun, the full-length *P. umbilicalis* cytochrome c_6_ linked to the `transit peptide from the` light-harvesting complex I chlorophyll a/b binding protein 6 (At3g54890) were used to generate an over-expression construct driven by the CaMV 35S promoter; B2-C6 in the vector pGWB2^42^. The recombinant plasmid B2-C6, was introduced into SBPase over-expressing tobacco cv. Samsun^3^ using *Agrobacterium tumefaciens* AGL1 via leaf-disc transformation^45^. Primary transformants (39) (T0 generation) were regenerated on MS medium containing kanamycin (100mg L^−1^), hygromycin (30 mg L^−1^) and augmentin (500 mg L^−1^). Plants expressing the integrated transgenes were screened using RT-PCR (data not shown).

In a similar fashion, the recombinant plasmids EC23083, and EC23028 were introduced into wild type tobacco (*Nicotiana tabacum*) cv Petit Havana, using *A. tumefaciens* strain LBA4404 via leaf-disc transformation^45^, and shoots were regenerated on MS medium containing, hygromycin (20 mg L^−1^) and cefotaxime (400 mg L^−1^). Hygromycin resistant primary transformants (T0 generation) with established root systems were transferred to soil and allowed to self-fertilize.

Between twelve and 60 independent lines were generated per construct and 3-4 lines were taken forward for full analysis. Control (CN) plants used in this study were a combined group of WT and null segregants from the transgenic lines, verified by PCR for non-integration of the transgene.

### Plant Growth

#### Controlled conditions

Wild-type tobacco plants and T1 progeny resulting from self-fertilization of transgenic plants were grown to seed in soil (Levington F2, Fisons, Ipswich, UK). Lines of interest were identified by qPCR. For the experiments in the Samsun cv. the null segregants were selected from transformed lines. For Petit Havana, the null segregants were selected from the S_B_C_6_ lines. For experimental study, T2-T4 and F1-F3 progeny seeds were germinated on soil in controlled environment chambers at an irradiance of 130 μmol photons m^−2^ s^−1^, 22°C, relative humidity of 60%, in a 16-h photoperiod. Plants were transferred to individual 8 cm pots and grown for two weeks at 130 μmol photons m^−2^ s^−1^, 22°C, relative humidity of 60%, in a 16-h photoperiod. Plants were transferred to 4 L pots and cultivated in a controlled environment glasshouse (16-h photoperiod, 25°C-30°C day/20°C night, and natural light supplemented under low light induced by cloud cover with high-pressure sodium light bulbs, giving 380-1000 μmol m^−2^ s^−1^ (high-light) from the pot level to the top of the plant, respectively). Positions of the plants were changed 3 times a week and watered regularly with a nutrient medium^46^. Plants were positioned such that at maturity, a near-to-closed canopy was achieved and the temperature range was maintained similar to the ambient external environment. Four leaf discs (0.8-cm diameter) were taken for FBPase activity. These disks were taken from the same areas of the leaf used for photosynthetic measurements, immediately plunged into liquid N_2_ and stored at −80°C.

#### Field

Plants were grown as described in Lopez-Calcagno *et al*^37^. The field site was situated at the University of Illinois Energy Farm (40.11°N, 88.21°W, Urbana, IL). Two different experimental designs were used in 2 different years.

2016: Plants were grown in rows, spaced 30 cm apart with the outer boundary being a wild-type border. The entire experiment was surrounded by two rows of wild-type borders. Plants were irrigated when required using rain towers. T2 seed was germinated and after 11 d were moved to individual pots (350 mL). The seedlings were grown in the glasshouse for further 15 d before being moved into the field, and allowed to grow in the field for 14 d before harvest.

2017: Two experiments were carried out two weeks apart. A blocks-within-rows design was used where one block holds one line of each of the five manipulations and each row has all lines. The central 20 plants of each block are divided into five rows of four plants per genotype. The 2017 Exp.1 contained controls (WT and null segregants), FBP/SBPase expressing lines (S_B_) and cytochrome *c*_6_ expressing lines (C_6_). The 2017 Exp. 2 contained controls (WT and null segregants), cytochrome *c*_6_ expressing lines (C_6_), and FBP/SBPase + cytochrome *c*_6_ expressing lines (S_B_C_6_). Seed was germinated and after 12 d moved to hydroponic trays (Trans-plant Tray GP009 6912 cells; Speedling Inc., Ruskin, FL), and grown in the glasshouse for 20 d before being moved to the field. The plants were allowed to grow in the field until flowering (approximately 30 d) before harvest.

The field was prepared in a similar fashion each year as described in Kromdijk *et al*.^47^. Light intensity (LI-quantum sensor; LI-COR) and air temperature (Model 109 temperature probe; Campbell ScientificInc, Logan, UT) were measured nearby on the same field site, and half-hourly averages were logged using a data logger (CR1000; Campbell Scientific).

### cDNA generation and RT-PCR

Total RNA was extracted from tobacco leaf disks (sampled from glasshouse grown plants and quickly frozen in liquid nitrogen) using the NucleoSpin^®^ RNA Plant Kit (Macherey-Nagel, Fisher Scientific, UK). cDNA was synthesized using 1 μg total RNA in 20 μl using the oligo-dT primer according to the protocol in the RevertAid Reverse Transcriptase kit (Fermentas, Life Sciences, UK). cDNA was diluted 1 in 4 to a final concentration of 12.5ng μL^−1^. For semi quantitative RT-PCR, 2 μL of RT reaction mixture (100 ng of RNA) in a total volume of 25 μL was used with DreamTaq DNA Polymerase (Thermo Fisher Scientific, UK) according to manufacturer’s recommendations. PCR products were fractionated on 1.0% agarose gels. For qPCR, the SensiFAST SYBR No-ROX Kit was used according to manufacturer’s recommendations (Bioline Reagents Ltd., London, UK). Primers used for semi quantitative RT-PCR can be seen in **Table 2.**

**Table 2;.**
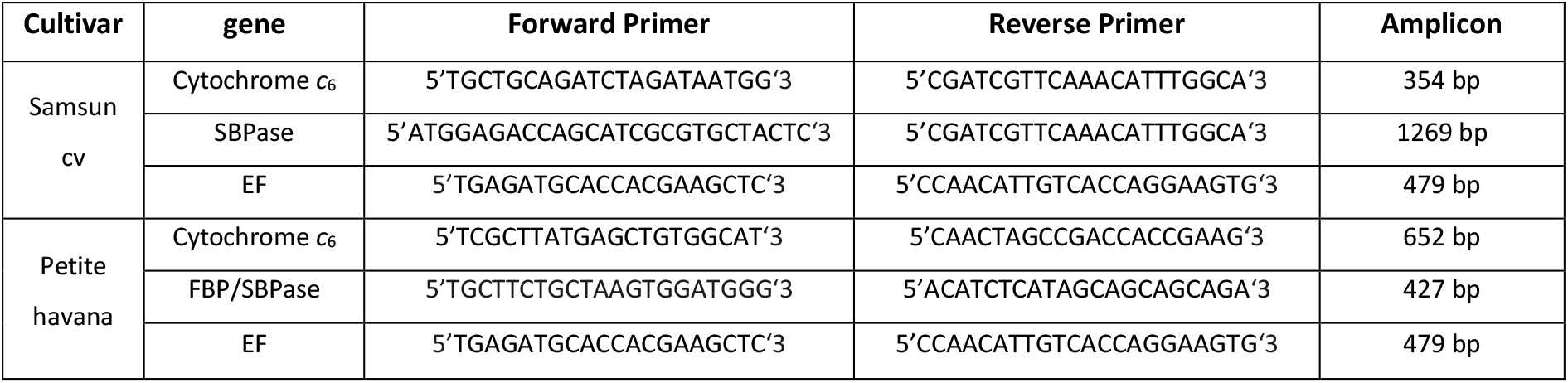
Primers used for semi-quantitative RT-PCR.

### Determination of FBPase Activity by Phosphate Release

FBPase activity was determined by phosphate release as described previously for SBPase with minor modifications^6^. Leaf discs were isolated from the same leaves and frozen in liquid nitrogen after photosynthesis measurements were completed. Leaf discs were ground to a fine powder in liquid nitrogen and immersed in extraction buffer (50 mM HEPES, pH8.2; 5 mM MgCl; 1 mM EDTA; 1 mM EGTA; 10% glycerol; 0.1% Triton X-100; 2 mM benzamidine; 2 mM aminocapronic acid; 0.5 mM phenylmethylsulfonylfluoride; 10 mM dithiothreitol), centrifuged 1 min at 14,000 xg, 4°C. The resulting supernatant (1 ml) was desalted through an NAP-10 column (Amersham) and stored in liquid nitrogen. The assay was carried out as descried in Simkin *et al*.^6^. In brief, 20 μl of extract was added to 80 μl of assay buffer (50 mM Tris, pH 8.2; 15 mM MgCl_2_; 1.5 mM EDTA; 10 mM DTT; 7.5 mM fructose-1,6-bisphosphate) and incubated at 25 °C for 30 min. The reaction was stopped by the addition of 50 μl of 1 M perchloric acid. 30 μl of samples or standards (PO^3-^_4_ 0.125 to 4 nmol) were incubated 30 min at room temperature following the addition of 300 μl of Biomol Green (Affiniti Research Products, Exeter, UK) and the A620 was measured using a microplate reader (VERSAmax, Molecular Devices, Sunnyvale, CA). Activities were normalized to transketolase activity^49^.

### Chlorophyll fluorescence imaging screening in seedlings

Chlorophyll fluorescence imaging was performed on 2-3 week-old tobacco seedlings grown in a controlled environment chamber at 130 μmol mol^−2^ s^−1^ and ambient (400 μmol mol^−1^) CO_2_. Chlorophyll fluorescence parameters were obtained using a chlorophyll fluorescence (CF) imaging system (Technologica, Colchester, UK^50,51^). The operating efficiency of photosystem two (PSII) photochemistry, *F*_q_’/*F*_m_’, was calculated from measurements of steady state fluorescence in the light (*F*’) and maximum fluorescence (*F*_m_’) following a saturating 800 ms pulse of 6300 μmol m^−2^ s^−1^ PPFD and using the following equation *F*_q_’/*F*_m_’ = (*F*_m_’-*F*’)/*F*_m_’. Images of *F*_q_’/*F*_m_’ were taken under stable PPFD of 600 μmol m^−2^ s^−1^ for Petite Havana and 650 μmol m^−2^ s^−1^ for Samsun^52–54^.

### Leaf Gas Exchange

Photosynthetic gas-exchange and chlorophyll fluorescence parameters were recorded using a portable infrared gas analyser (LI-COR 6400; LI-COR, Lincoln, NE, USA) with a 6400-40 fluorometer head unit. Unless stated otherwise, all measurements were taken with LI-COR 6400 cuvette. For plants grown in the glasshouse conditions were maintained at a CO_2_ concentration, leaf temperature and vapour pressure deficit (VPD) of 400 μmol mol^−1^, 25 °C and 1 ± 0.2 kPa respectively. The chamber conditions for plants grown under field conditions had a CO_2_ concentration of 400 μmol mol^−1^, block temperature was set to 2 °C above ambient temperature (ambient air temperature was measure before each curve) and VPD was maintained as close to 1 kPa as feasible possible.

### *A*/*C*_i_ response curves (Photosynthetic capacity)

The response of net photosynthesis (*A*) to intracellular CO_2_ concentration (*C*_i_) was measured at a saturating light intensity of 2000 μmol mol^−2^ s^−1^. Illumination was provided by a red-blue light source attached to the leaf cuvette. Measurements of *A* were started at ambient CO_2_ concentration (C_a_) of 400 μmol mol^−1^, before C_a_ was decreased step-wise to a lowest concentration of 50 μmol mol^−1^ and then increased step-wise to an upper concentration of 2000 μmol mol^−1^. To calculate the maximum saturated CO_2_ assimilation rate (*A*_max_), maximum carboxylation rate (*Vc*_max_) and maximum electron transport flow (*J*_max_), the C3 photosynthesis model^55^ was fitted to the *A*/*C*_i_ data using a spreadsheet provided by Sharkey *et al*.^56^. Additionally, chlorophyll fluorescence parameters including PSII operating efficiency (*F*_q_’/*F*_m_’) and the coefficient of photochemical quenching (*q*P), mathematically identical to the PSII efficiency factor (*F*_q_’/*F*_v_’) were recorded at each point.

### *A*/*Q* response curves

Photosynthesis as a function of light (*A*/*Q* response curves) was measured under the same cuvette conditions as the *A*/*C*_i_ curves mentioned above. Leaves were initially stabilized at saturating irradiance of 2200 to μmol m^−2^ s^−1^, after which *A* and *g*_s_ were measured at the following light levels: 2000, 1650, 1300, 1000, 750, 500, 400, 300, 200, 150, 100, 50 and 0 μmol m^−2^ s^−1^). Measurements were recorded after *A* reached a new steady state (1-3 min) and before *g*_s_ changed to the new light levels. Values of *A* and *g*_s_ were used to estimate intrinsic water-use efficiency (*iWUE* = *A*/*g*_s_)

### Statistical Analysis

All statistical analyses were done using Sys-stat, University of Essex, UK, and R (https://www.r-project.org/). For harvest data, seedling chlorophyll imaging and enzyme activities, analysis of variance and Post hoc Tukey test were done. For gas exchange curves, data were compared by linear mixed model analysis using lmer function and type III anova^57^. Significant differences between manipulations were identified using contrasts analysis (lsmeans package).

## Acknowledgments

This study was supported by the Realising Improved Photosynthetic Efficiency (RIPE) initiative awarded to C.A.R by University of Illinois, USA. RIPE was possible through support from the Bill & Melinda Gates Foundation, DFID and FFAR, grant OPP1172157. This work was also supported by the Biotechnology and Biological Sciences Research Council (BBSRC) grant BB/J004138/1. We would like to thank Jack Matthews (University of Essex) for help with data analysis, Elena A. Pelech (University of Illinois) and Sunitha Subramaniam (University of Essex) for help with plant growth, Phillip A. Davey (University of Essex) and Richard Gossen (University of Helsinki) for help with gas exchange and David Drag, Ben Harbaugh and the Ort lab (University of Illinois) for support with the field trials.

## Author contributions

P.E.L.C and A.J.S. generated transgenic plants. P.E.L.C, A.J.S, K.L.B. and S.J.F. performed molecular and biochemical experiments. P.E.L.C, A.J.S and K.L.B carried out plant phenotypic and growth analysis and performed gas exchange measurement. A.J.S and S.J.F performed enzyme assays on selected lines; all authors carried out data analysis on their respective contributions; C.A.R and T.L designed and supervised the research; P.E.L.C., A.J.S and C.A.R wrote the manuscript, P.E.L.C, K.L.B. and A.J.S contributed equally to the completion of this work.

